# Sensitivity and specificity of a Bayesian single trial analysis for time varying neural signals

**DOI:** 10.1101/690958

**Authors:** Jeff T. Mohl, Valeria C. Caruso, Surya T. Tokdar, J. M. Groh

## Abstract

We recently reported the existence of fluctuations in neural signals that may permit neurons to code multiple simultaneous stimuli sequentially across time^1^. This required deploying a novel statistical approach to permit investigation of neural activity at the scale of individual trials. Here we present tests using synthetic data to assess the sensitivity and specificity of this analysis. We fabricated datasets to match each of several potential response patterns derived from single-stimulus response distributions. In particular, we simulated dual stimulus trial spike counts that reflected fluctuating *mixtures* of the single stimulus spike counts, stable *intermediate* averages, *single* stimulus winner-take-all, or response distributions that were *outside* the range defined by the single stimulus responses (such as summation or suppression). We then assessed how well the analysis recovered the correct response pattern as a function of the number of simulated trials and the difference between the simulated responses to each “stimulus” alone. We found excellent recovery of the mixture, intermediate, and outside categories (>97% correct), and good recovery of the single/winner-take-all category (>90% correct) when the number of trials was >20 and the single-stimulus response rates were 50Hz and 20Hz respectively. Both larger numbers of trials and greater separation between the single stimulus firing rates improved categorization accuracy. These results provide a benchmark, and guidelines for data collection, for use of this method to investigate coding of multiple items at the individual-trial time scale.

## Introduction

We recently showed that when multiple stimuli are present, some neurons exhibit activity patterns that fluctuate between those evoked by each stimulus alone^1^. This dynamic code could allow the representation of all stimuli within the same population of neurons. Such fluctuations may be a widespread phenomenon in the brain, but would be overlooked using conventional analysis methods that investigate mean activity pooled across trials. Of particular interest are cases in which the time-and-trial-pooled responses evoked by multiple stimuli appear to reflect the average of the responses to each stimulus presented in isolation. This phenomenon, known as divisive normalization^2^, has been observed in visual brain areas such as V1 and MT^3^ as well as other sensory and cognitive domains^4–9^. However, such responses could either reflect a true averaging of the two stimuli/conditions, producing a consistent stable intermediate level of firing on each trial, or could reflect a dynamic code that flexibly shifts between the individual stimuli across trials.

To evaluate neural responses on a single trial basis, the novel statistical approach introduced in Caruso et al. (2018) characterizes the distribution of spike counts elicited in response to two simultaneous stimuli using Bayesian inference. Here we provide a general assessment of the sensitivity and specificity of that approach by simulating known neural responses as benchmark cases. In particular, we investigate how the analysis performs as we parametrically varied the data sample size (number of trials), the mean firing rate of responses, and the difference between spike counts across conditions.

We demonstrate that our approach accurately categorizes synthetic neural data into expected categories. The robustness of the results depends heavily on sample size, as well as on firing rate differences between the two single cue conditions. Importantly, the model performs very well under reasonable experimental values (20 trials per condition, 60% firing rate change between conditions). Finally, we show that that the model gracefully handles datasets that do not exactly match any of the tested hypotheses. These results demonstrate the viability of the analysis method and provide constraints for interpretation of actual neural data.

## Experimental Rationale and Procedures

### Neural encoding patterns to be assessed

For simplicity, our approach focused on the case of two simultaneously presented stimuli (dual stimuli) but can be extended to a larger number of stimuli. We consider an experimental setup in which a neuron’s response is recorded in three interleaved conditions: in the presence of a single stimulus “A”, a single stimulus “B”, or both stimuli A and B (“AB”). We considered four possible response distributions to dual stimuli, in relation to the distributions observed when only one stimulus is present (Figure 1).

**Figure 1.**
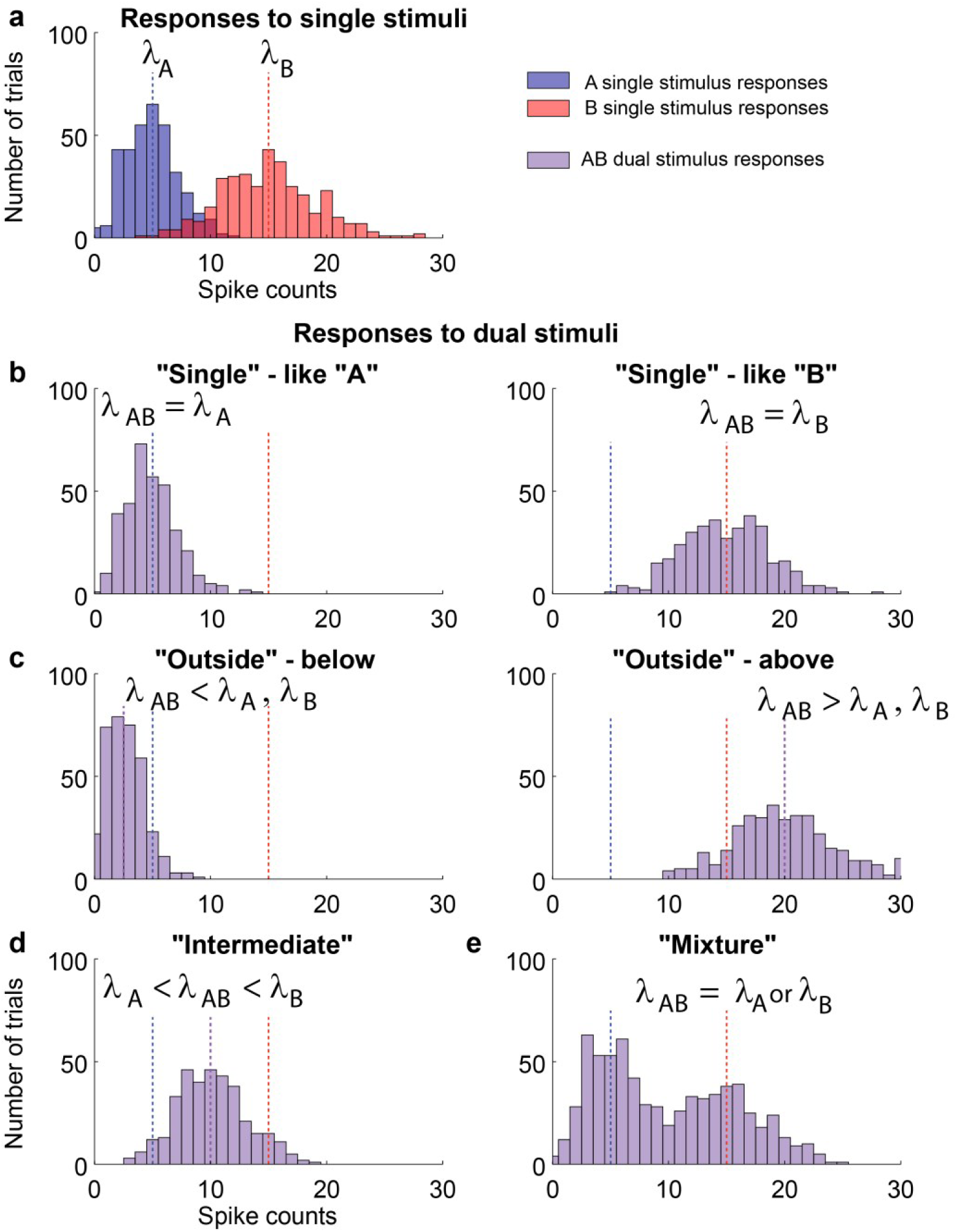
Four possible response patterns to dual stimuli trials, in relation to the responses observed to the component stimuli when presented individually. (**a**) Single stimulus trials were modeled as evoking spike counts distributed according to a Poisson process with rates λ_A_, blue, or λ_B_, red. (**b**) Responses on dual stimulus trials follow a Poisson λ_AB_ matching either λ_A_, left, or λ_B_, right. (**c**) The Poisson rate λ_AB_ on dual stimulus trials is less than (left) or greater than (right) those observed in single stimulus trials. In these simulations, λ_AB_ was set to 0.5* λ_A_ (left) or λ_A_+λ_B_ (right). (**d**) Spike counts derived from a Poisson process with a rate λ_AB_ between λ_A_ and λ_B_. (**e**) Spike counts drawn from a mixture of two Poissons with rates λ_A_ and λ_B_.

1. Neurons might respond to only one of the stimuli, and do so consistently (i.e respond to the same one) across trials. One way this could occur would be if only one stimulus is located in a neuron’s receptive field, but it might also apply when both stimuli are in the receptive field (sometimes referred to as a winner-take-all encoding). We label this possibility “single”.
2. The responses to dual stimuli might be greater than the maximum or less than the minimum of the single-stimulus responses. This category includes enhancement/summation, as well as suppression of the response to one stimulus by another. We refer to this as “outside”.
3. The responses to dual stimuli are a *consistent* weighted average of the responses evoked by each stimulus alone. Here, the dual stimulus responses are between the bounds set by the two single stimulus responses, and cluster around a stable intermediate value. We refer to this case as “intermediate”.
4. The responses to dual stimuli may fluctuate such that on each trial the neuron appears to be responding to only one of the two stimuli. We term this possibility “mixture” because it reflects a mixture of two distributions of A and B stimulus responses. This is analogous to a winner-take-all except that the neuron is switching across trials rather than encoding the same stimulus each trial. Like the “intermediate” category, there could be a weighting factor such that a higher proportion of trials favor one stimulus over the other.

### Model construction, Bayesian model comparison, and synthetic data

These four possibilities can be formalized on the basis of how the spike distributions on combined stimulus trials AB compare to those observed when only A or B are presented alone. If A and B elicit spike counts according to Poisson distributions *Poi*(*λ*_*A*_) and *Poi*(*λ*_*B*_), then we can ask which of four competing hypotheses best describe the spike counts observed on combined AB trials:

a. Single: *F* = *Poi*(*λ*) for either *λ* = *λ*_*A*_ or *λ* = *λ*_*B*_, with *λ* constant across trials
b. Outside: *F* = *Poi*(*λ*) for some unknown *λ* ∉ (min(*λ*_*A*_, *λ*_*B*_), max(*λ*_*A*_, *λ*_*B*_))
c. Intermediate: *F* = *Poi*(*λ*) for some unknown *λ* ∈ (min(*λ*_*A*_, *λ*_*B*_), max(*λ*_*A*_, *λ*_*B*_))
d. Mixture: *F* = *α* · *Poi*(*λ*_*A*_) + (1 − *α*) · *Poi*(*λ*_*B*_) for some unknown *α* ∈ (0,1)

The plausibility of each of these models was determined by computing the posterior probabilities of each model given the data, with a default Jeffreys’ prior^10^ on each of the model specific rate (*λ*) parameters and on the mixing probability parameter (*α*). Each model was given a uniform prior probability (1/4) and posterior model probabilities were calculated by computation of relevant intrinsic Bayes factors^11^ (see appendix S1 for a thorough description of the models and model selection strategy).

To evaluate the sensitivity and specificity of this method, we built synthetic neuronal spiking datasets to match each of the four potential encoding strategies tested by the model. Consistent with our previous study^1^, we focused on response patterns that could be modeled as deriving from Poisson distributions. In principle, the approach could be extended to other forms of response distributions, but this is beyond the scope of this work.

Data files were generated as spike times drawn using an independent Poisson point process sampled at 1 ms Intervals, with constant mean firing rate for 1000 ms (Figure 2a-c). For A and B (single stimulus) trials, Poisson rates were assigned a priori to reflect a range of realistic firing rates for a single neuron presented with different stimuli. AB (combined stimulus) trials for each dataset were generated based on the chosen A and B firing rates and in a manner consistent with one of the four potential hypotheses. For the “single” hypothesis the AB data were generated using a single Poisson with rate λ_AB_ equal to the highest of the component rates, i.e. max(λ_A_, λ_B_). For the “outside” hypothesis, the rate λ_AB_ was set 20% higher than max(λ_A_, λ_B_). For the “intermediate” hypothesis, λ_AB_ was equal to the mean of A and B rates λ_AB_ = 0.5 (λ_A_) + 0.5(λ_B_). For the across trial switching (mixture) model, the data were generated using the same Poisson process, but each trial was randomly chosen to be drawn from either poi(λ_A_) or poi(λ_B_) with equal probability. This results in a dataset for which the across trial average firing rate is equal to the average of the λ_A_ and λ_B_ rates, but individually each trial is better described as deriving from either the λ_A_ or λ_B_ response distributions. Note that it is nearly impossible to tell by visual inspection of a raster plot when a neuron has such a mixed response pattern, even when the trials are sorted as they are in Figure 2c, but the pattern becomes more evident in histograms of the trial-wise spike counts (Figure 2d).

**Figure 2.**
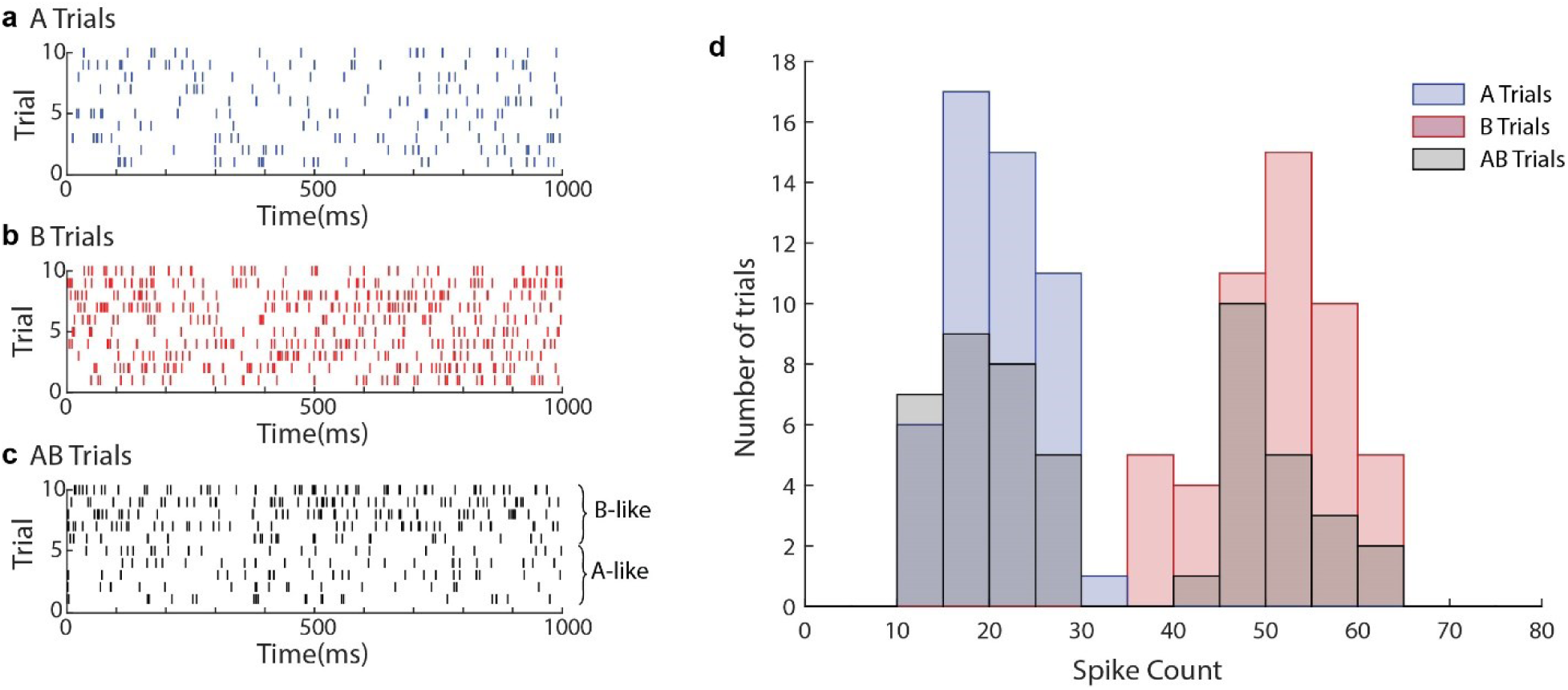
One example synthetic dataset. (**a-c**) Raster plots for a synthetic dataset built to match the across trial switching hypothesis, where blue rows (**a**) are single A trials, red rows (**b**) are single B trials, and black rows (**c**) are AB trials. AB trials are drawn randomly to match either A or B rates and are sorted so that B-like rates are towards the top of the raster. Even with sorting, this pattern is challenging to see with the naked eye, highlighting the need for analytical methods. (**d**) Whole trial spike count histogram for 50 repetitions of A, B, and AB trials. From this plot, the bimodality of AB trials for the switching condition is more apparent.

Multiple datasets were generated using this strategy in order to test the power and reliability of the analysis under plausible experimental conditions. These datasets varied both the number of trials per condition (5-50 trials per condition) and the firing rates of the A and B conditions (from 1-100 Hz, with relative separation between A and B rates of 33-80% of maximum rate). Individual triplet pairs were generated under each of these conditions, analogous to running 100 individual cells through the analysis. This set of parameters was used for all conditions tested, including datasets constructed to not exactly match any of the hypotheses, discussed in the final section of the results.

### Code and data availability

Code specific to this paper can be found on GitHub at https://github.com/jmohl/mplx_tests, (archived DOI: 10.5281/zenodo.3508536) which includes the code used to generate synthetic data for this manuscript as well as all code needed to perform the neural mixture analysis. The exact synthetic data files used to generate plots are available upon request. Source code and documentation for the Neural Mixture Model available at https://github.com/tokdarstat/Neural-Multiplexing.

## Results

### Neural Mixture Model accurately characterizes synthetic data built to match hypotheses

The desired analysis outcome is for the output to match the input. That is, data explicitly generated to match the single hypothesis should be correctly labeled as “single”, data generated as “outside” should be labelled “outside”, etc. Figure 3 illustrates that this is largely the case. The series of simulations shown here involved 20 A trials simulated with λ_A_ = 20 Hz, 20 B trials with λ_B_ = 50 Hz, and 20 AB trials generated according to the various methods specified above. The “mixture” and “intermediate” categories perform exceptionally well, with 100/100 “mixture” and 99/100 “intermediate” datasets labeled correctly with >95% confidence (dark black bars). This distinction is critical, as these two conditions would produce exactly the same mean rate when averaging across trials, making them indistinguishable using typical neural analysis strategies which average across trials in order to reduce noise. “Single” and “outside” datasets were also correctly labeled in the majority of cases (90/100 and 97/100, respectively), although these hypotheses are not the focus of our analysis as they can be differentiated more easily using simpler statistical methods.

**Figure 3.**
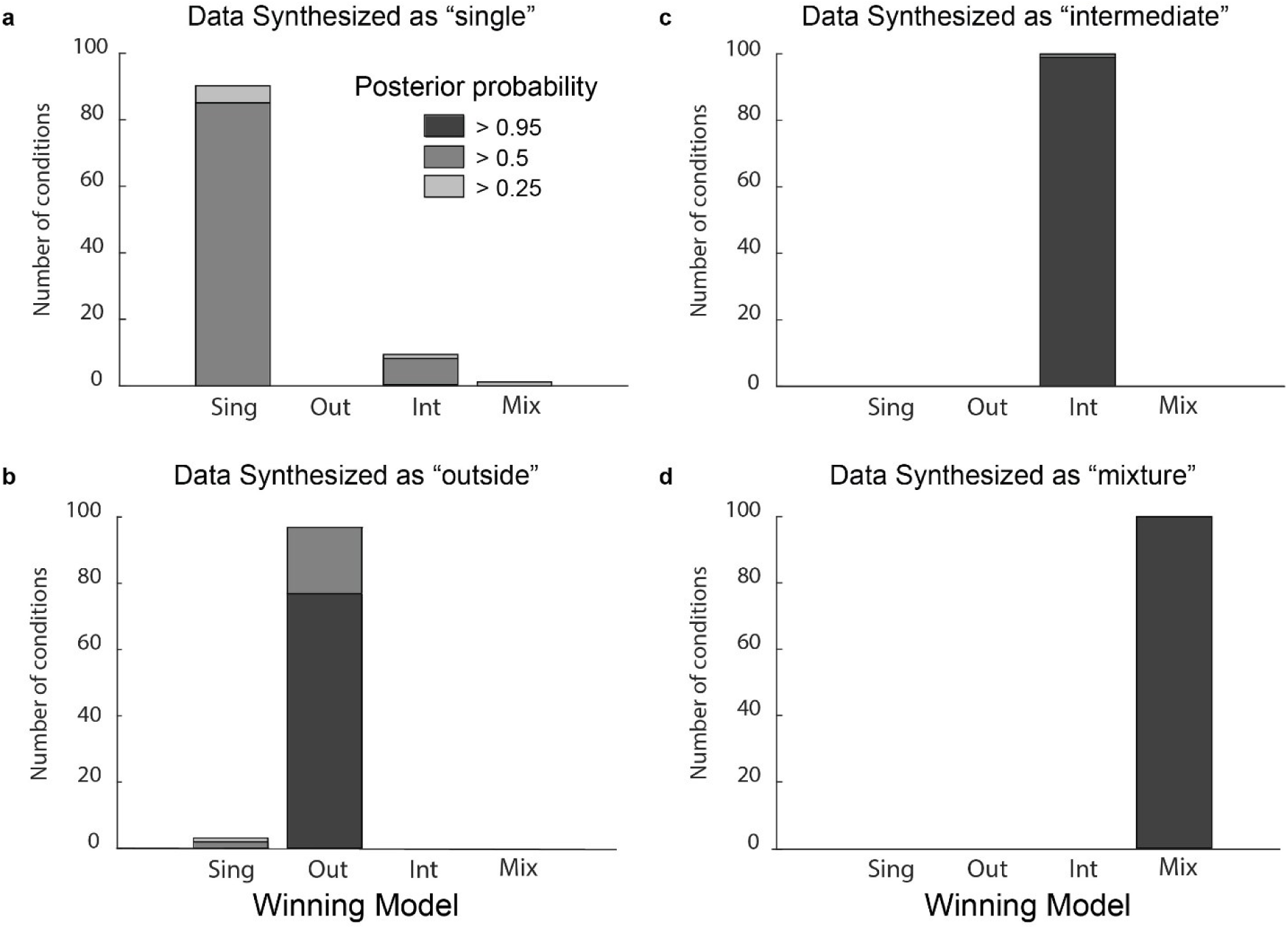
The analysis method correctly categorizes synthetic datasets created to match each model. The shading of the bars indicates the posterior probability with which each individual run of synthetic data (n=100) is assigned to a given category. Of particular interest is the very strong separation between intermediate and mixture datasets, as this discrimination is not possible when considering only firing rates averaged across trials. Parameters used for this figure: λ_A_ = 20Hz, λ_B_=50 Hz, number of trials in each run = 20 per stimulus condition (60 overall).

Although the category “single” was correctly identified as the best model for the dataset simulated under the “single” hypothesis 90% of the time, the posterior probability or confidence level did not reach the 95% level observed for the other models. This is due to the narrow definition of this category: response rates on AB trials must be indistinguishable from those occurring on either the A or B trials. All other categories include a range of possibilities which admits this hypothesis as a boundary case (i.e. a weighted average with the weight for A set to 1). Therefore, these models are all competitive in explaining data that is generated to match the “single” case, which explains the low posterior probability of this model. For this reason, it is better to consider the “single” category as reflective of a null hypothesis, where there is no interaction at all between stimuli.

### Dependence on number of trials and difference between A and B response distributions

The accuracy of this characterization depended on both the amount of data and the difference between the response distributions on A and B trials. The dependence on the number of trials is best appreciated when considering similar A and B response distributions, such as *λ*_*A*_ *=* 20 vs *λ*_*B*_ *=* 30Hz as shown in Figure 4a (right), which depicts the average posterior probability value for the correct model as a function of the number of trials. Even with this modest separation between the A and B response patterns, increasing the number of trials per condition allowed the analysis to better characterize the underlying rates, and therefore better discriminate between competing hypotheses. “Single”, “Intermediate” and “Mixture” had average posterior probability values >0.3 for N=5 trials, but performance improves steadily to average posterior probability values of >0.75 for N=50 trials. When response distributions were moderately separated, *λ*_*A*_ *=* 20 vs *λ*_*B*_ *=* 50Hz (the same separation used in Figure 3), performance rose more rapidly for all models except “single”. At N=5, posterior probability values range from 0.4 for “single” to 0.8 for “mixture”. At N=30, posterior probability values equaled approximately 1 for “mixture”, “intermediate” and “outside”. Further increasing the firing rate separation to *λ*_*A*_ *=* 20 vs *λ*_*B*_ *=* 100Hz resulted in very high posterior probabilities of >0.95 even at N=5 for “mixture” and “intermediate”; this level was achieved for “outside” at N=10.

**Figure 4.**
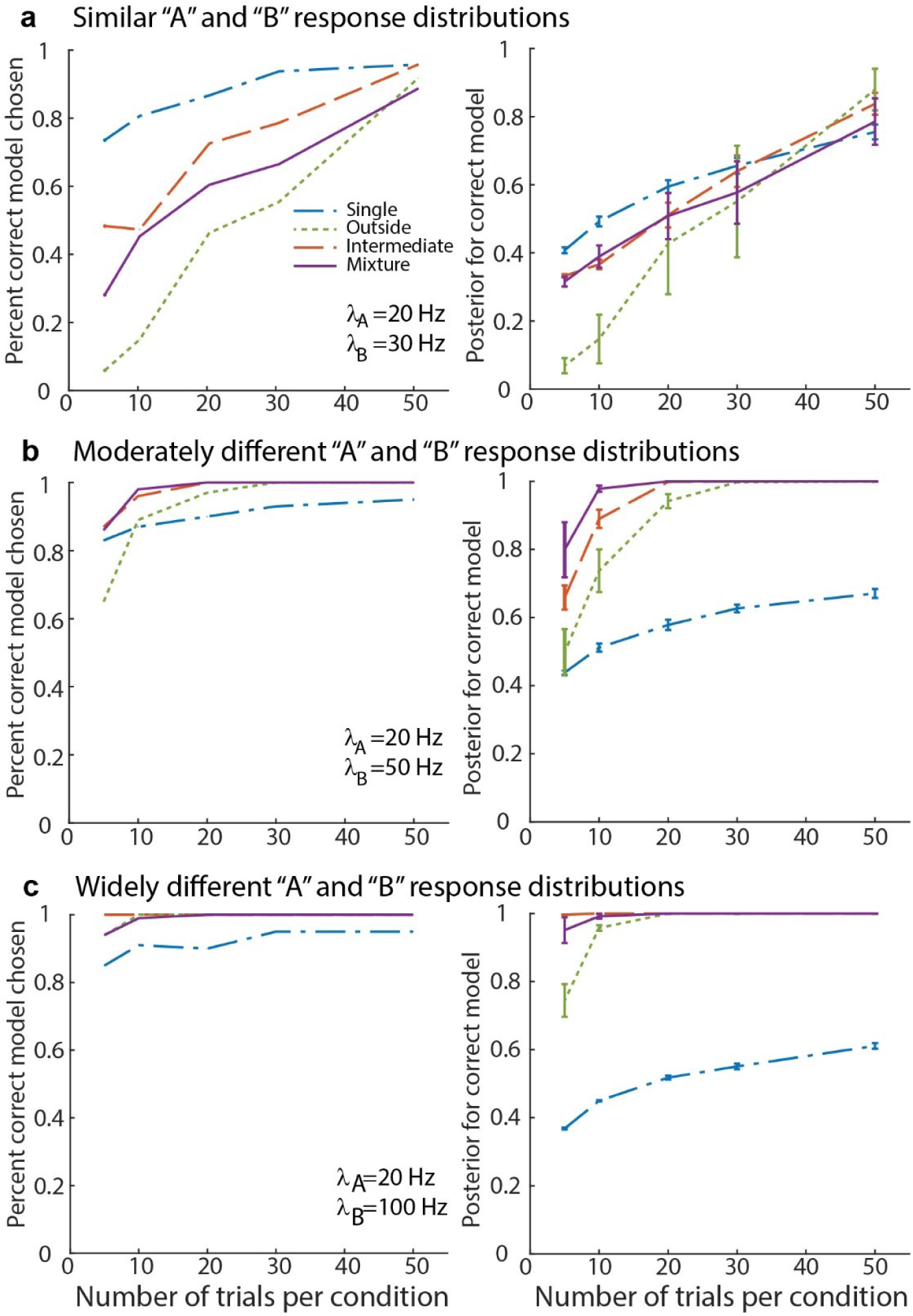
Increasing number of trials or separability of conditions improves accuracy of model comparison. (**a**) Left, percent of triplets which were correctly categorized, split by dataset type, for increasing number of trials per conditions; λ_A_ = 20Hz, λ_B_=30 Hz. Right, mean and variance of posterior probability for correct model across triplets (**b** & **c**) same as in **a** but with λ_B_ set to 50z and 100 Hz respectively. Fewer trials are needed when responses are very different between A and B trials.

These figures give a rough sense of the sensitivity of our analysis, demonstrating that the analysis becomes more reliable as more trials per condition are added until reaching asymptote around 30 trials/condition for a 50 Hz vs 20 Hz comparison (Figure 4b). Similarly, increasing the difference in spike count between A and B conditions also improves specificity in the analysis, allowing for accurate characterization with as few as 5 trials (Figure 4c). Although these data were constructed under ideal conditions (the data perfectly matches one of the tested hypotheses), they can be used as a guide for how much data should be collected in order to obtain satisfactory results in a real dataset.

The above results give a detailed view of how each model performs across a realistic range of firing rates for neural recordings from primate sensory cortices and sub-cortical areas, for which we initially designed this method. We next sought to determine whether the analysis effectively extended into datasets with much lower maximum firing rates. To address this question we performed the analysis on data for a wide range of average firing rate values (5 to 100 Hz) for two fixed amounts of relative separation between A and B rates (40% and 80% of maximum rate) and characterized the prediction accuracy under each model (Figure 5). We found that average firing rate affected the accuracy of the model, with lower average firing rate conditions resulting in worse performance than higher firing rates. As expected, a larger relative separation between A and B responses (analogous to having stronger neuronal preference for one or the other condition) resulted in significantly better performance, even for low firing rate conditions. However, even when considering the larger separation value of 80%, datasets with a maximum firing rate of <15 Hz barely reached 95% accuracy with 50 trials per condition. These results suggest that datasets with very low average firing rates (less than ∼15 Hz for the most preferred response) may not be resolvable using this analysis method.

**Figure 5.**
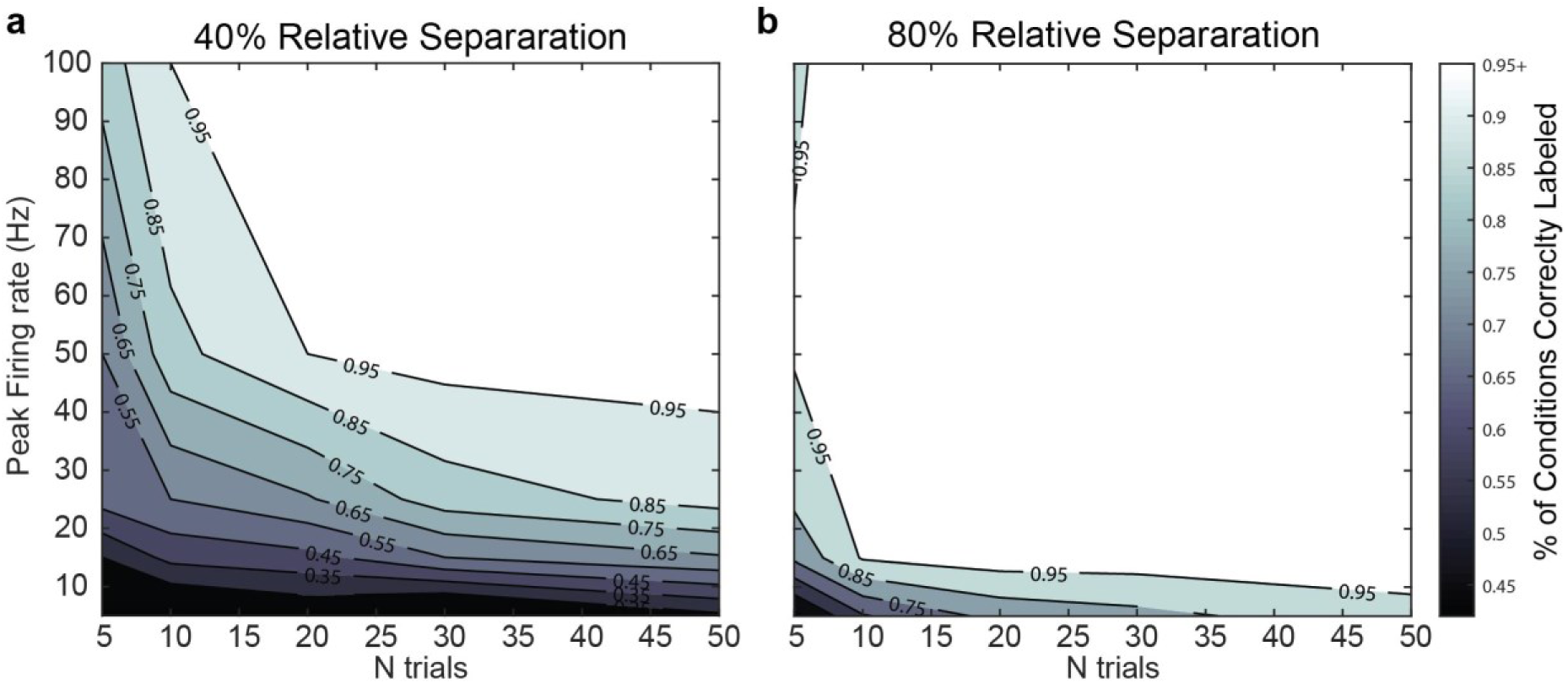
Model prediction accuracy depends on both number of trials and average firing rate. (**a**) Model classification accuracy collapsed across all four models for datasets generated for a fixed relative separation between A and B responses of 40% of the peak firing rate. Shading shows the percentage of conditions correctly categorized with interpolated phase transitions. (**b**) Same as in **a**, but for a relative separation of 80% of the peak rate.

### Model results are informative even for datasets that do not perfectly match hypotheses

Though this modeling strategy was meant to test between discrete hypotheses, it is unlikely that real neural signals perfectly and uniquely match any one of these scenarios. Therefore, it is important that the analysis accurately reflect deviations from exact hypothesis matches. Here, we consider two potential deviations from the circumstances considered above: weighted averaging of A and B stimuli and incomplete switching between A and B rates.

For weighted averaging datasets the AB trials were generated as in the “intermediate” condition above, except that the AB rate was set to be closer to the A rate than the B rate: *λ*_*AB*_ = 0.75 * *λ*_*A*_ + 0.25 * *λ*_*B*_. Because the model returns both a classification and a posterior probability (reflecting the model’s confidence in that classification), we expected that this type of dataset will result in a spread across single and average classifications, but with lower confidence in this assessment. This is indeed the case, as the analysis returned primarily the intermediate category, with some single winners, but with much lower posterior probabilities than the well matched datasets (Figure 6a, compare with Figure 3c).

**Figure 6.**
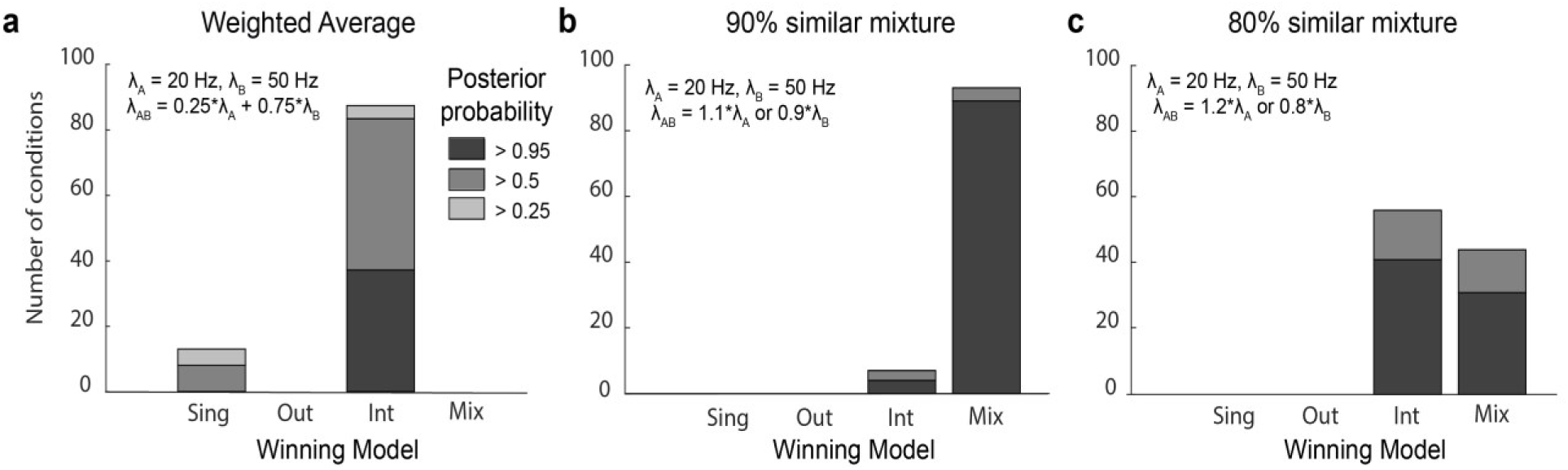
Datasets which do not exactly match the canonical hypotheses are descriptively categorized by model comparison. (**a**) A dataset generated to reflect weighted average of A and B stimuli is weakly categorized as intermediate with some triplets categorized as singles. (**b**) Mixture trials generated to alternate between values shifted 10% from the true A and B firing rates are primarily categorized as mixtures. (**c**) Mixture trials with rates shifted 20% from the true A and B rates are categorized as either mixture or average with equal probability, consistent with the fact that these trials would be much more difficult to discriminate from the true averaging hypothesis. Parameters used for this figure: λ_B_ = 50Hz, λ_A_=20 Hz, number of trials = 20.

We also tested a form of incomplete switching, where stimuli show strong fluctuations but did not quite exactly match the A and B rates. These datasets were generated using the same strategy as the mixture datasets described above, except that A-like or B-like trials were generated with a slightly shifted mean firing rate. Multiple degrees of similarity were tested, but two (80% and 90% similarity) are presented here. From these, the analysis accurately described a 90% switch as mixture (Figure 6b), but around 80% similarity it began to interpret many datasets as average (Figure 6c). This highlights a natural limitation that should be expected in the data, as continuing to reduce the similarity would eventually result in a condition that was indistinguishable from true averaging. However, these results demonstrate the high specificity of our analysis for the mixture category, enforcing a strong definition of mixture (literally switching between rates closely matched to the A and B rates, rather than any amount of fluctuation).

## Discussion

There is broad interest in understanding the nature and significance of firing patterns in the brain. It is well known that such firing patterns are variable in the face of identical, highly controlled experimental conditions (such as the presentation of the same stimulus in the same context). While many studies have viewed this variability as deleterious “noise” that if unsolved would undermine the ability of the brain to perform its essential tasks^12–16^, we and others have sought explanations under the possibility that certain forms of such variation may contribute in a positive fashion to brain function^1,17–24^. In particular, we have successfully modeled variation in whole-trial spike counts to multiple stimuli as being drawn from the observed distributions of spike counts to those same stimuli when presented individually^1^.

Here, we benchmarked this analysis on synthetic datasets to provide insight into the sensitivity and specificity of our analysis method as a function of trial counts and the separation between the distributions of spike counts elicited by the component individual stimuli. When response separations are large, e.g. the mean rate for one stimulus condition is 5X the rate for the other, the analysis method can successfully distinguish among the 4 competing hypotheses with as few as n=5 trials for each condition (n=15 overall). Smaller response separations can be compensated for by collecting more trials to achieve similarly good results. Finally, even when the conditions do not exactly match the assumptions, such as if the component response rates in the “mixture” condition do not precisely match those observed when the single stimuli were presented individually, correct classifications greatly outnumber incorrect ones. Critically, the analysis is conservative against the “mixture” hypothesis in these cases, demonstrating that data which is best fit by this model is truly fluctuating between the responses to single stimuli at the single trial level.

These results suggest that the analysis tested here are suitable for many electrophysiological datasets which match the A, B, AB format. Datasets which high peak firing rates (∼50 Hz) and an average response difference between conditions of approximately 40% (relative to peak rate) can reach over 95% categorization accuracy with as few as 20 trials. Higher peak firing rates, larger separation between neural response, or a greater number of trials all improve the accuracy of our analysis. Conversely, datasets with low peak firing rates (∼15 Hz) are likely to produce only weak results even with a large number of trials (which will be reflected in low model posterior probabilities). Practically, this means that our analysis is particularly well suited for recordings in primate sensory or motor brain regions with the pronounced tuning and firing rate changes required to differentiate responses at the single trial level.

A situation not tested here is the case in which the response distributions are not derived from Poisson distributions. In our previous work^1^ we excluded conditions in which the responses to individual stimuli did not satisfactorily resemble Poisson distributions in order to ensure that our model assumptions were not violated, but this has several downsides. First, it is difficult to have confidence in the success of this exclusion criterion: failing to reject the Poisson assumption is not the same as confirming its validity. Second, a considerable amount of data is excluded in this fashion (as much as 25-50%, depending on dataset, before even considering other exclusion criteria). Finally, there is significant evidence in the literature that spike counts in many brain areas are more variable than would be suggested by a Poisson distribution^15,25–29^. For all of these reasons, it will be important to both test the model with data sets that violate this assumption and extend the analysis method to include other response distributions such as negative binomials.

The data presented here reflect conditions where two “stimuli” are presented at the same time, but this analysis could in principle be extended to combinations of three or more response patterns. We have previously shown that responses to multiple auditory stimuli in the primate inferior colliculus are often well described by mixtures of single stimulus responses^1^, but little is known about how this type of code changes as more stimuli are added. An extension of this analysis into more complex mixtures of multiple different responses may help bring clarity to this question, and more work is needed to determine whether this phenomenon is general or limited to two stimulus cases.

Given the broad interest in both noise as a potential limitation on neural representations and in divisive normalization as an elemental computation in sensory processing – with recent suggestions that this process may be impaired in conditions such as autism^30–32^ – it will be increasingly important to develop additional methods which can probe neural codes at the individual trial level^33–35^. The tools described in the present paper represent an important step towards uncovering fluctuating patterns in neural activity that may permit greater amounts of information to be encoded in the spike trains of individual and populations of neurons.

