## Supplemental Methods for "Sensitivity and specificity of a Bayesian single trial analysis for time varying neural signals"

### 1 Introduction

Here we record in detail the modeling strategy used and specifics of the model selection procedure, the results of which are reported in the main text.

### 2 Model

For each experimental condition $e \in \{\mathrm{A}, \mathrm{B}, \mathrm{AB}\}$, let $Y_j^e$, $j = 1, \ldots, n_e$ denote the spike counts from all $n_e$ trials run under the condition. We model

1. $Y_j^{\mathrm{A}} \sim Poi(\lambda_{\mathrm{A}})$, $Y_j^{\mathrm{B}} \sim Poi(\lambda_{\mathrm{B}})$ for some unknown $\lambda_{\mathrm{A}}, \lambda_{\mathrm{B}} > 0$; and,
2. $Y_j^{\mathrm{AB}} \sim F$ with four competing hypotheses describing $F$
   (a) Mixture: $F = \alpha \cdot Poi(\lambda_{\mathrm{A}}) + (1 - \alpha) \cdot Poi(\lambda_{\mathrm{B}})$ for some unknown $\alpha \in (0, 1)$
   (b) Intermediate: $F = Poi(\lambda)$ for some unknown $\lambda \in (\min(\lambda_{\mathrm{A}}, \lambda_{\mathrm{B}}), \max(\lambda_{\mathrm{A}}, \lambda_{\mathrm{B}}))$
   (c) Outside: $F = Poi(\lambda)$ for some unknown $\lambda \notin [\min(\lambda_{\mathrm{A}}, \lambda_{\mathrm{B}}), \max(\lambda_{\mathrm{A}}, \lambda_{\mathrm{B}})]$
   (d) Single: $F = Poi(\lambda)$ for either $\lambda = \lambda_{\mathrm{A}}$ or $\lambda = \lambda_{\mathrm{B}}$, with the exact situation being unknown *a priori*.

### 3 Method

#### 3.1 Bayesian testing

We carry out statistical testing between the set of hypotheses $H$ listed above by adopting a Bayesian perspective. A prior probability $p_h$ is assigned to each hypothesis $h \in H$, with $p_h > 0$ and $\sum_{h \in H} p_h = 1$. Let the observed data be denoted $Y = (Y_j^e : 1 \leq j \leq n_e, e \in \{\mathrm{A}, \mathrm{B}, \mathrm{AB}\})$. Each hypothesis $h$ gives rise to a model for $Y$ which can be written generically as

$$Y \sim f_h(y \mid \theta_h), \theta_h \in \Theta_h$$

where $\theta_h$ captures all unknown parameters under the hypothesis. The modeling process is completed by assuming a prior distribution $\pi_h(\theta_h)$ on the parameter space $\Theta_h$ to reflect prior information and beliefs about the uncertainty about $\theta_h$. Inference about $\theta_h$ is then drawn based on the uncertainty quantified by the resulting posterior distribution $\pi_h(\theta_h \mid Y) = \pi_h(\theta_h) f_h(Y \mid \theta_h)/f_h(Y)$ over $\Theta_h$ where the normalizing constant

$$f_h(Y) := \int_{\Theta_h} f_h(Y \mid \theta_h)\pi_h(\theta_h)d\theta_h$$

is recognized as the *marginal likelihood score* for hypothesis $h$ given the observed data. Inference about the relative merits of the competing hypotheses is then drawn based on the posterior hypothesis probabilities

$$p_h(Y) = \frac{p_h f_h(Y)}{\sum_{h' \in H} p_{h'} f_{h'}(Y)}, h \in H, \tag{1}$$

which capture the post-data certainties about the competing hypotheses.

### 3.2 Prior specification

Because the four competing hypotheses are only about the distribution of AB trial counts, and do not differ in their description of A and B trial count distributions, we adopt a common prior for the parameters pertaining to these latter distributions. Specifically, we take $\lambda_{\text{A}} \sim \mathit{Gam}(a, b)$ and $\lambda_{\text{B}} \sim \mathit{Gam}(a, b)$ for each of the four models.

For the Mixture hypothesis, the remaining model parameter is the mixing proportion $\alpha \in (0, 1)$. We assign it a beta prior: $\alpha \sim \mathit{Be}(c_1, c_2)$. For the Intermediate hypothesis, given $\lambda_{\text{A}}$ and $\lambda_{\text{B}}$, the remaining parameter $\lambda$ is assigned a conditional gamma prior $\mathit{Gam}(a, b)$ truncated to the interval $(\min(\lambda_{\text{A}}, \lambda_{\text{B}}), \max(\lambda_{\text{A}}, \lambda_{\text{B}}))$. Similarly, for the Outside hypothesis, we take the conditional prior on $\lambda$ given $\lambda_{\text{A}}, \lambda_{\text{B}}$ as the $\mathit{Gam}(a, b)$ distribution truncated to $(0, \infty) \cap [\min(\lambda_{\text{A}}, \lambda_{\text{B}}), \max(\lambda_{\text{A}}, \lambda_b)]^c$. In both these cases, the same $a, b$ values are used as for the prior distributions for $\lambda_{\text{A}}$ and $\lambda_{\text{B}}$.

### 3.3 Computation

Computation of marginal likelihood scores $f_h(Y)$ is generally a complex task in Bayesian inference and require customized approaches to numerically evaluate the integration. Our prior choices and the low dimensionality of the parameter spaces associated with all the hypotheses make the task slightly easier for our problem. However, each hypothesis demands a different strategy to perform the integration and we give enough details below so that an enterprising student can implement these strategies from scratch.

Before getting into the details, we note a particular simplification that can be made to the expression of $p_h(Y)$ in (1) thanks to the assumption of a common prior distribution on $\lambda_{\text{A}}$ and $\lambda_{\text{B}}$ across all four hypotheses. Write

$Y = (Y^{\mathrm{A}}, Y^{\mathrm{B}}, Y^{\mathrm{AB}})$ where each $Y^e$ denotes the data corresponding to experimental condition $e \in \{\mathrm{A}, \mathrm{B}, \mathrm{AB}\}$. Then we can write

$$p_h(Y) = \frac{p_h \tilde{f}_h(Y^{\mathrm{AB}} \mid Y^{\mathrm{A}}, Y^{\mathrm{B}})}{\sum_{h' \in H} p_{h'} \tilde{f}_{h'}(Y^{\mathrm{AB}} \mid Y^{\mathrm{A}}, Y^{\mathrm{B}})} \tag{2}$$

where

$$\begin{aligned} \tilde{f}_h(Y^{\mathrm{AB}} \mid Y^{\mathrm{A}}, Y^{\mathrm{B}}) = \int \Big\{ & \tilde{f}_h(Y^{\mathrm{AB}} \mid \tilde{\theta}_h, \lambda_{\mathrm{A}}, \lambda_{\mathrm{B}}) \pi_h(\tilde{\theta}_h \mid \lambda_{\mathrm{A}}, \lambda_{\mathrm{B}}) \times \\ & \pi(\lambda_{\mathrm{A}} \mid Y^{\mathrm{A}}) \pi(\lambda^{\mathrm{B}} \mid Y^{\mathrm{B}}) \Big\} d\tilde{\theta}_h d\lambda_{\mathrm{A}} d\lambda_{\mathrm{B}} \end{aligned}$$

with $\tilde{f}_h$ denoting the probability mass function of $Y^{\mathrm{AB}}$ under model $h$ and $\tilde{\theta}_h$ denoting the remaining parameters of the model. Notice that

$$\pi(\lambda_{\mathrm{A}} \mid Y^{\mathrm{A}}) = Gam(a + S_{\mathrm{A}}, b + n_{\mathrm{A}}), \quad \pi(\lambda_{\mathrm{B}} \mid Y^{\mathrm{B}}) = Gam(a + S_{\mathrm{B}}, b + n_{\mathrm{B}}) \tag{3}$$

where $S_{\mathrm{A}} = \sum_{j=1}^{n_{\mathrm{A}}} Y_j^{\mathrm{A}}$ and $S_{\mathrm{B}} = \sum_{j=1}^{n_{\mathrm{B}}} Y_j^{\mathrm{B}}$.

A particular integration operation that shows up repeatedly in the following calculations stems from the well-know Poisson-Gamma conjugacy. For a vector of non-negative integers $y = (y_1, \ldots, y_n)$ and positive real numbers $\alpha, \beta$, we define the quantity

$$\begin{aligned} g(y; \alpha, \beta) &:= \int \prod_{j=1}^{n} Poi(y_j \mid \lambda) Gam(\lambda \mid \alpha, \beta) d\lambda & (4) \\ &= \frac{\Gamma(\alpha + S(y))}{\Gamma(\alpha)} \frac{\beta^\alpha}{(\beta + n)^{\alpha + S(y)}} \frac{1}{\prod_{j=1}^{n} y_j!} & (5) \end{aligned}$$

where $S(y) = \sum_{j=1}^{n} y_j$. This quantity could be easily evaluated using any standard mathematics or statistics software. But we do want caution the user of numerical overflow problems as the gamma function grows super-exponentially in its argument. It is best to carry out the calculation of $g(y; \alpha, \beta)$ in the logarithmic scale, such as using the `lgamma()` function in the R software platform.

#### 3.3.1 Computation for "Single" hypothesis

Leveraging the Poisson-Gamma conjugacy, one can directly calculate

$$\begin{aligned} \tilde{f}(Y^{\mathrm{AB}} \mid Y^{\mathrm{A}}, Y^{\mathrm{B}}) &= \frac{1}{2} \Big[ \int \prod_{j=1}^{n_{\mathrm{AB}}} Poi(Y_j^{\mathrm{AB}} \mid \lambda_{\mathrm{A}}) \pi(\lambda_{\mathrm{A}} \mid Y^{\mathrm{A}}) d\lambda_{\mathrm{A}} \\ &\qquad + \int \prod_{j=1}^{n_{\mathrm{AB}}} Poi(Y_j^{\mathrm{AB}} \mid \lambda_{\mathrm{B}}) \pi(\lambda_{\mathrm{B}} \mid Y^{\mathrm{B}}) d\lambda_{\mathrm{B}} \Big] \\ &= \frac{1}{2} \Big[ g(Y^{\mathrm{AB}}; a + S_{\mathrm{A}}, b + n_{\mathrm{A}}) + g(Y^{\mathrm{AB}}; a + S_{\mathrm{B}}, b + n_{\mathrm{B}}) \Big]. \end{aligned}$$

The above calculation is done by assuming that the total prior probability of the Single hypothesis is split equally (*a priori*) between its two sub-hypotheses. A more conservative variation of this would be to report the maximum of the two numbers $g(Y^{\mathrm{AB}}; a + S_{\mathrm{A}}, b + n_{\mathrm{A}})$ and $g(Y^{\mathrm{AB}}; a + S_{\mathrm{B}}, b + n_{\mathrm{B}})$, giving full weight to the sub-hypothesis that explains the data better. Such selective representation of strongest sub-hypothesis has been used in the literature (Berger and Guglielmi, 2001).

#### 3.3.2 Computation for "Mixture" hypothesis

For this hypothesis, the remaining parameter is $\tilde{\theta} = \alpha$ with $\pi(\alpha \mid \lambda_{\mathrm{A}}, \lambda_{\mathrm{B}}) = Be(c_1, c_2)$ and

$$\tilde{f}(Y^{\mathrm{AB}} \mid \alpha, \lambda_{\mathrm{A}}, \lambda_{\mathrm{B}}) = \prod_{j=1}^{n_{\mathrm{AB}}} \{\alpha \cdot Poi(Y_j^{\mathrm{AB}} \mid \lambda_{\mathrm{A}}) + (1-\alpha) \cdot Poi(Y_j^{\mathrm{AB}} \mid \lambda_{\mathrm{B}})\}. \quad (6)$$

This form of $\tilde{f}$ is difficult to work with directly because when the product of the sum is expanded, it results in too many summands; $2^{n_{\mathrm{AB}}}$ many to be precise. Instead, a commonly adopted strategy in dealing with mixture model computation is to rewrite the model by introducing latent (unobserved) variables $Z_j \in \{\mathrm{A}, \mathrm{B}\}$, $j = 1, \ldots, n_{\mathrm{AB}}$ that indicate which of the two Poisson components observation $j$ came from. By considering,

$$Z_j \sim Discrete(\{\mathrm{A}, \mathrm{B}\}; (\alpha, 1-\alpha)); \quad Y_j \mid (Z_j = c) \sim Poi(\lambda_c); \quad j = 1, \ldots, n_{\mathrm{AB}}$$

we recover the same joint distribution $\tilde{f}$ for $Y^{\mathrm{AB}}$ as in (6).

One may write $\tilde{f}(Y^{\mathrm{AB}} \mid Y^{\mathrm{A}}, Y^{\mathrm{B}}) = \int \tilde{f}(Y^{\mathrm{AB}} \mid Y^{\mathrm{A}}, Y^{\mathrm{B}}, Z) p(Z) dZ$ where $p(Z)$ denotes the joint distribution on $Z$ under the model (and the integral actually is a sum over a discrete space): $p(Z) = \int p(Z \mid \alpha)\pi(\alpha) d\alpha = B(c_1 + \#\{Z_j = \mathrm{A}\}, c_2 + \#\{Z_j = \mathrm{B}\})/B(c_1, c_2)$ where $B(\cdot, \cdot)$ is the beta function. The integral can be numerically approximated by importance sampling Monte Carlo as follows. Let $q(Z)$ denote any distribution on the space of $Z$. Then with $Z^m, m = 1, \ldots, M$, denoting a large sample of independent draws of $Z$ from $q(Z)$, one has

$$\tilde{f}(Y^{\mathrm{AB}} \mid Y^{\mathrm{A}}, Y^{\mathrm{B}}) \approx \frac{1}{M} \sum_{m=1}^{M} \tilde{f}(Y^{\mathrm{AB}} \mid Y^{\mathrm{A}}, Y^{\mathrm{B}}, Z = Z^m) \frac{p(Z = Z^m)}{q(Z = Z^m)}$$

by the strong law of large numbers. The quality of this Monte Carlo approximation is improved by choosing an *importance distribution* $q(Z)$ that closely resembles the posterior distribution $p(Z \mid Y^{\mathrm{AB}}, Y^{\mathrm{A}}, Y^b) \propto \tilde{f}(Y^{\mathrm{AB}} \mid Y^{\mathrm{A}}, Y^{\mathrm{B}}, Z)p(Z)$; see (Tokdar and Kass, 2010) for more details. For our purposes, a good and convenient choice is a $q(Z)$ under which $Z_j \sim Discrete(\{\mathrm{A}, \mathrm{B}\}, (\bar{\alpha}_j, 1 - \bar{\alpha}_j))$,

$j = 1, \ldots, n$, where

$$\bar{\alpha}_j = \frac{\int Poi(Y_j^{\mathrm{AB}} \mid \lambda_{\mathrm{A}})\pi(\lambda_{\mathrm{A}} \mid Y^{\mathrm{A}})d\lambda_{\mathrm{A}}}{\int Poi(Y_j^{\mathrm{AB}} \mid \lambda_{\mathrm{A}})\pi(\lambda_{\mathrm{A}} \mid Y^{\mathrm{A}})d\lambda_{\mathrm{A}} + \int Poi(Y_j^{\mathrm{AB}} \mid \lambda_{\mathrm{B}})\pi(\lambda_{\mathrm{B}} \mid Y^{\mathrm{B}})d\lambda_{\mathrm{B}}},$$
$$= \frac{g(Y_j^{\mathrm{AB}}; a + S_{\mathrm{A}}, b + n_{\mathrm{A}})}{g(Y_j^{\mathrm{AB}}; a + S_{\mathrm{A}}, b + n_{\mathrm{A}}) + g(Y_j^{\mathrm{AB}}; a + S_{\mathrm{B}}, b + n_{\mathrm{B}})},$$

which calculates the probability of classifying trial $j$ as having come from condition A, under equal prior odds.

Therefore, to carry out the above importance sampling Monte Carlo, it is sufficient that we evaluate $\tilde{f}(Y^{\mathrm{AB}} \mid Y^{\mathrm{A}}, Y^{\mathrm{B}}, Z)$. But this can be computed efficiently as

$$\tilde{f}(Y^{\mathrm{AB}} \mid Y^{\mathrm{A}}, Y^{\mathrm{B}}, Z) = \int \tilde{f}(Y^{\mathrm{AB}} \mid \lambda^{\mathrm{A}}, \lambda^{\mathrm{B}}, Z)\pi(\lambda_{\mathrm{A}} \mid Y^{\mathrm{A}})\pi(\lambda^{\mathrm{B}} \mid Y^{\mathrm{B}})d\lambda_{\mathrm{A}}d\lambda_{\mathrm{B}}$$
$$= \left\{ \int \prod_{j:Z_j = \mathrm{A}} Poi(Y_j^{\mathrm{AB}} \mid \lambda_{\mathrm{A}})\pi(\lambda_{\mathrm{A}} \mid Y^{\mathrm{A}})d\lambda_{\mathrm{A}} \right\}$$
$$\times \left\{ \int \prod_{j:Z_j = \mathrm{B}} Poi(Y_j^{\mathrm{AB}} \mid \lambda_{\mathrm{B}})\pi(\lambda_{\mathrm{B}} \mid Y^{\mathrm{B}})d\lambda_{\mathrm{B}} \right\}$$
$$= g(\{Y_j^{\mathrm{AB}} : Z_j = \mathrm{A}\}; a + S_{\mathrm{A}}, b + n_{\mathrm{A}}) \cdot g(\{Y_j^{\mathrm{AB}} : Z_j = \mathrm{B}\}; a + S_{\mathrm{B}}, b + n_{\mathrm{B}})$$

by using the Poisson-Gamma conjugacy.

#### 3.3.3 Computation for the "Intermediate" hypothesis

For the Intermediate hypothesis, one can use a straight Monte Carlo average to approximate $\tilde{f}(Y^{\mathrm{AB}} \mid Y^{\mathrm{A}}, Y^{\mathrm{B}})$ as

$$\tilde{f}(Y^{\mathrm{AB}} \mid Y^{\mathrm{A}}, y^{\mathrm{B}})$$
$$= \int \left\{ \int \tilde{f}(Y^{\mathrm{AB}} \mid \lambda)\pi(\lambda \mid \lambda_{\mathrm{A}}, \lambda_{\mathrm{B}})d\lambda \right\} \pi(\lambda_{\mathrm{A}} \mid Y^{\mathrm{A}})\pi(\lambda_{\mathrm{B}} \mid Y^{\mathrm{B}})d\lambda_{\mathrm{A}}d\lambda_{\mathrm{B}}$$
$$\approx \frac{1}{M} \sum_{m=1}^{M} \tilde{f}(Y^{\mathrm{AB}} \mid \lambda_{\mathrm{A}} = \lambda_{\mathrm{A}}^{m}, \lambda_{\mathrm{B}} = \lambda_{\mathrm{B}}^{m})$$

where $(\lambda_A^m, \lambda_B^m)$, $m = 1, \ldots, M$, are independent draws from $\pi(\lambda_A \mid Y^A) \times \pi(\lambda_B \mid Y^B)$ and, with $\underline{\lambda} = \min(\lambda_A, \lambda_B)$, $\overline{\lambda} = \max(\lambda_A, \lambda_B)$, $S_{AB} = \sum_{j=1}^{n_{AB}} Y_j^{AB}$,

$$\begin{aligned}
\tilde{f}(Y^{AB} \mid \lambda_A, \lambda_B) &= \int \tilde{f}(Y^{AB} \mid \lambda)\pi(\lambda \mid \lambda_A, \lambda_B)d\lambda \\
&= \frac{\int_{\underline{\lambda}}^{\overline{\lambda}} \prod_{j=1}^{n} Poi(Y_j^{AB} \mid \lambda)\lambda^{a-1}e^{-b\lambda}d\lambda}{\int_{\underline{\lambda}}^{\overline{\lambda}} \lambda^{a-1}e^{-b\lambda}d\lambda} \\
&= \frac{\int_{\underline{\lambda}}^{\overline{\lambda}} \lambda^{a+S_{AB}-1}e^{-(b+n_{AB})\lambda}d\lambda}{\{\prod_{j=1}^{n} Y_j^{AB}!\} \int_{\underline{\lambda}}^{\overline{\lambda}} \lambda^{a-1}e^{-b\lambda}d\lambda} \\
&= g(Y^{AB}; a, b) \times \frac{F_{a+S_{AB},b+n}(\overline{\lambda}) - F_{a+S_{AB},b+n}(\underline{\lambda})}{F_{a,b}(\overline{\lambda}) - F_{a,b}(\underline{\lambda})}
\end{aligned}$$

where $F_{\alpha,\beta}(x)$ is used to denote the cumulative distribution function of $Gam(\alpha, \beta)$.

#### 3.3.4 Computation for "Outside" hypothesis

Here the computation is done exactly as in the Intermediate hypothesis case, except for the following calculation which accounts for the fact that the conditional prior on $\lambda$ given $\lambda_A$, $\lambda_B$ is $Gam(a, b)$ truncated to the complement of the interval $(\underline{\lambda}, \overline{\lambda})$:

$$\tilde{f}(Y^{AB} \mid \lambda_A, \lambda_B) = g(Y^{AB}; a, b) \times \frac{1 - \{F_{a+S_{AB},b+n}(\overline{\lambda}) - F_{a+S_{AB},b+n}(\underline{\lambda})\}}{1 - \{F_{a,b}(\overline{\lambda}) - F_{a,b}(\underline{\lambda})\}}.$$

### 3.4 Additional considerations for non-informative priors

An actual implementation of the above testing framework requires choosing the hyper-parameters $a, b, c_1, c_2$, all positive valued real numbers. As with any Bayesian analysis, the results will have some dependence on the choice of these hyper-parameters. While expert knowledge about the model animal, brain region and sensory/cognitive task might help to choose reasonable values of these parameters, it may also be desirable to use some *default* values that encode minimal prior information about the model parameters.

One such approach is to use non-informative priors arising from Jeffreys' work. For the prior on the mixing proportion $\alpha$, the Jeffreys prior is $Be(1/2, 1/2)$ which corresponds to our choice with $c_1 = c_2 = 1/2$. The Jeffreys' prior for estimating the mean of a Poisson distribution is the improper density function $\pi(\mu) \propto 1/\sqrt{\mu}$, $\mu > 0$, which matches, in a limiting sense, our choice of $Gam(a, b)$ with $a = 1/2$ and $b = 0$. This is because the posterior distribution for the Poisson mean under a $Gam(1/2, \beta)$ prior converges to the posterior distribution under the Jeffreys' prior as $\beta \to 0$.

However, such a limiting property does not hold for the marginal likelihood score! In fact, this score is not even well defined under the Jeffreys' prior, since prior density function is defined only up to a multiplicative constant. Also note that the quantity $g(y; \alpha, \beta) \to 0$ as $\beta \to 0$, and, hence working with a small but non-zero $b$ is not an option either, since the resulting marginal likelihood scores for the Intermediate and the Outside hypotheses can be made arbitrarily small by choosing an arbitrarily small $b$.

Such anomalies can be effectively addressed by following the intrinsic Bayes factor approach of Berger and Pericchi (1996). The Bayes factor between two hypotheses $h$ and $h'$ is defined as the ratio of the marginal likelihood scores $B_{h,h'}(Y) = f_h(Y)/f_{h'}(Y)$. Notice that

$$\frac{p_h(Y)}{p_{h'}(Y)} = \frac{p_h}{p_{h'}} \times B_{h,h'}(Y), \tag{7}$$

that is the posterior odds between the two hypotheses depends on the data $Y$ only through the Bayes factor. The intrinsic Bayes factor adjustment works for the case where data $Y$ consists of a collection of observations $(Y_1, \ldots, Y_n)$ which, under each hypothesis $h$, are independently distributed with their distributions depending on a parameter $\theta_h \in \Theta_h$, with a prior distribution $\pi_h(\theta_h)$ chosen on $\Theta_h$.

When one or both of $\pi_h(\theta_h)$ and $\pi_{h'}(\theta_{h'})$ are improper, defined only up to an arbitrary scaling factor, Berger and Pericchi (1996) recommend replacing them with proper distributions $\pi_h^\ell(\theta_h) = \pi_h(\theta_h \mid Y_\ell)$ and $\pi_{h'}^\ell(\theta_{h'}) = \pi_{h'}(\theta_{h'} \mid Y_\ell)$ obtained by calculating the posterior distribution given a small fraction of the data $Y_\ell = (Y_j : j \in \ell)$, for subset $\ell \subset \{1, \ldots, n\}$ called a training set. A minimal training set is chosen, so that least amount of data is expended in this step of correcting for the impropriety of the prior distributions. Next, one calculates the marginal likelihood scores based on the new priors $\pi_h^\ell(\theta_h)$ and $\pi_{h'}^\ell(\theta_{h'})$, but using only the remaining part of the data $Y \setminus Y_\ell$. The resulting Bayes factor, which depends on the choice of the training set, but does not depend on any arbitrary scaling of the original priors, can be expressed as $B_{h,h'}^\ell(Y) = B_{h,h'}(Y)/B_{h,h'}(Y_\ell)$. To avoid the the effect of the arbitrary choice of the training set, one calculates the intrinsic Bayes factor $B_{h,h'}^*(Y)$ which is an average of $B_{h,h'}^\ell$ across all minimal training sets $\ell$.

The final averaging could be an arithmetic, geometric or harmonic mean of the training set adjusted Bayes factors. We adopt the geometric mean approach, because it generalizes nicely to the case where one has more than two hypotheses to compare. The geometric mean intrinsic Bayes factor preserves the transitivity property that $B_{h,h''}^* = B_{h,h'}^* \times B_{h',h''}^*$ and conforms to (7) with $B$ replaced with $B^*$, for every pair of hypotheses $h, h' \in H$. Furthermore, the geometric mean intrinsic Bayes factor approach can be viewed as a direct adjustment to the marginal likelihood score $B_{h,h'}^*(Y) = f_h^*(Y)/f_{h'}^*(Y)$, where the corresponding intrinsic marginal likelihood score $f_h^*(Y)$ is defined as the geometric mean of $f_h(Y)/f_h(Y_\ell)$ across all minimal training sets $\ell$.

For our four hypotheses, both Outside and Intermediate have improper priors when $b = 0$. For either hypothesis, a single observation is enough to give a proper posterior and hence the minimal training set size is one. Therefore the intrinsic marginal likelihood score adjustment for any or our models is achieved as:

$$\tilde{f}_h^*(Y^{\mathrm{AB}} \mid Y^{\mathrm{A}}, Y^{\mathrm{B}}) = \frac{\tilde{f}_h(Y^{\mathrm{AB}} \mid Y^{\mathrm{A}}, Y^{\mathrm{B}})}{\left[\prod_{\ell=1}^{n_{\mathrm{AB}}} \tilde{f}_h(Y_\ell^{\mathrm{AB}} \mid Y^{\mathrm{A}}, Y^{\mathrm{B}})\right]^{1/n_{\mathrm{AB}}}} \tag{8}$$

where one uses the formulas derived above for $\tilde{f}_h(Y^{\mathrm{AB}} \mid Y^{\mathrm{A}}, Y^{\mathrm{B}})$ with $b \approx 0$. In our implementation we use $b = 10^{-5}$. The adjustments to the Single, Intermediate and Outside hypotheses are straightforward. For the Mixture model, one does not need to run an importance sampling Monte Carlo to calculate the denominator in (8). Instead, since for each $\ell \in \{1, \ldots, n\}$ the corresponding $Z_\ell$ has only two possibilities $\{\mathrm{A}, \mathrm{B}\}$ a full enumeration can be done to express the denominator as $(c_1 + c_2)^{-1}[\prod_{\ell=1}^{n_{\mathrm{AB}}}(c_1 g(Y_\ell^{\mathrm{AB}}; a + S_{\mathrm{A}}, b + n_{\mathrm{A}}) + c_2 g(Y_\ell^{\mathrm{AB}}; a + S_{\mathrm{B}}, b + n_{\mathrm{B}})]^{1/n_{\mathrm{AB}}}$.
